# Comparison of undirected frequency-domain connectivity measures for cerebro-peripheral analysis

**DOI:** 10.1101/2021.06.22.449393

**Authors:** Joachim Gross, Daniel S. Kluger, Omid Abbasi, Nikolas Chalas, Nadine Steingräber, Christoph Daube, Jan-Mathijs Schoffelen

**Affiliations:** Institute for Biomagnetism and Biosignal Analysis, University of Münster, Münster, Germany; Otto-Creutzfeldt-Center for Cognitive and Behavioral Neuroscience, University of Münster, Münster, Germany; Centre for Cognitive Neuroimaging, University of Glasgow, Glasgow, UK; Radboud University, Donders Institute for Brain, Cognition and Behaviour, Nijmegen, NL

**Author notes:** **Corresponding author**: Daniel S. Kluger, University of Münster, Institute for Biomagnetism and Biosignal Analysis, Malmedyweg 15, 48149 Münster, Germany.

**Keywords:** cerebro-peripheral connectivity, spectral analysis, functional connectivity, phase coupling

## Abstract

Analyses of cerebro-peripheral connectivity aim to quantify ongoing coupling between brain activity (measured by MEG/EEG) and peripheral signals such as muscle activity, continuous speech, or physiological rhythms (such as pupil dilation or respiration). Due to the distinct rhythmicity of these signals, undirected connectivity is typically assessed in the frequency domain. This leaves the investigator with two critical choices, namely a) the appropriate measure for spectral estimation (i.e., the transformation into the frequency domain) and b) the actual connectivity measure. As there is no consensus regarding best practice, a wide variety of methods has been applied. Here we systematically compare combinations of six standard spectral estimation methods (comprising fast Fourier and continuous wavelet transformation, bandpass filtering, and short-time Fourier transformation) and six connectivity measures (phase-locking value, Gaussian-Copula mutual information, Rayleigh test, weighted pairwise phase consistency, magnitude squared coherence, and entropy). We provide performance measures of each combination for simulated data (with precise control over true connectivity), a single-subject set of real MEG data, and a full group analysis of real MEG data. Our results show that, overall, wppc and gcmi tend to outperform other connectivity measures, while entropy was the only measure sensitive to bimodal deviations from a uniform phase distribution. For group analysis, choosing the appropriate spectral estimation method appeared to be more critical than the connectivity measure. We discuss practical implications (sampling rate, SNR, computation time, and data length) and aim to provide recommendations tailored to particular research questions.

## 1. Introduction

The analysis of cerebro-peripheral connectivity has recently gained significant interest. This analysis approach is typically based on two recordings with high temporal resolution, namely MEG/EEG recordings of brain activity (Baillet, 2017; Gross, 2019) and a peripheral signal sampled at the same rate. A prominent early application of cerebro-peripheral connectivity was the investigation of connectivity between brain and muscle activity (Salenius et al., 1997), which has led to important insights into the role of neural rhythms in physiological and pathological motor control (Bourguignon et al., 2019, 2017; Schnitzler & Gross, 2005; Schoffelen et al., 2005). More recently, this type of analysis has also proven useful for studying continuous speech processing due to the fact that brain signals are temporally synchronised to the speech envelope (Gross et al., 2013b; Lakatos et al., 2019; Meyer et al., 2019; Obleser & Kayser, 2019; Zoefel, 2018). More generally, cerebro-peripheral connectivity can be studied to elucidate the ongoing coupling between any peripherally recorded signal and brain activity (Gross, 2019; Park et al., 2014; Rebollo et al., 2018) and even modulations of such connectivity measures as a function of a secondary peripheral signal such as respiration (Kluger & Gross, 2020). Examples for relevant peripheral signals are eye movements, pupil size, heart beat, respiration, speech, movement or muscle activity, skin conductance or temperature, and blood pressure. Some of these signals (such as respiration, heart beat, speech, tremor) are distinctively rhythmic, thus favouring analysis in the spectral domain. However, there is no consensus in the literature on the best methodological approach to quantify cerebro-peripheral connectivity in the spectral domain. Instead, a large variety of methods has been used. In practice, spectral cerebro-peripheral connectivity analysis consists of two steps that can each be conducted in several ways: First, spectral estimation is performed where time series are transformed into the frequency domain (as complex-valued numbers). Spectral estimation is most often performed by using Fourier transformation, wavelet transformation, or bandpass-filtering (Bruns, 2004; Gross, 2014; Le Van Quyen and Bragin, 2007). In a second step, connectivity measures can be estimated. Again, a large number of methods have been suggested (Bastos and Schoffelen, 2015; Marzetti et al., 2019) and some of them have been compared in previous studies (David et al., 2004; Kreuz et al., 2007; Quian Quiroga et al., 2002). It is noteworthy that MEG/EEG connectivity is often discussed in the context of cerebro-cerebral connectivity, i.e. connectivity between different brain areas. This brings about complications that are absent in the case of cerebro-peripheral connectivity. Most importantly, estimation of non-invasive MEG/EEG time series from two regions of interest in the brain is never perfect and leads to leakage effects that contaminate the connectivity estimate (Schoffelen & Gross, 2009). This is typically circumvented using connectivity measures that exclude common zero-lag components in both time series (such as imaginary coherence). In the case of cerebro-peripheral connectivity, the estimation of time series in the brain is still not optimal but the second signal is a peripheral recording that does not share any spurious signal components with the brain signal that result from imperfect source reconstruction. Therefore, analyses of cerebro-peripheral connectivity do not require connectivity measures to exclude shared zero-lag signals.

Depending on the differences of multiple methods for spectral decomposition and estimation of effect size, the investigator’s choice could affect the results of the analysis. Here, we aim to investigate the sensitivity of cerebro-peripheral connectivity analysis to the choice of spectral estimation and connectivity measures. We realise that such an investigation depends on the signals that are used and on the implementation of the spectral estimation and connectivity methods. Therefore, we cannot expect to provide authoritative guidance on the ‘optimal’ analysis approach that generalises to all possible applications. Still, we can expect to learn lessons that could be valuable to the community in the planning of similar studies and the analysis of cerebro-peripheral data.

A second contribution is to make our analysis scripts publicly available on GitHub (https://github.com/IBiomag/) so that a similar comparison can be performed for different simulated or real data and different methods can be added and evaluated.

Since we anticipated non-trivial interactions between different spectral estimation methods and different connectivity measures, we analysed all combinations of a set of six standard spectral estimation methods (comprising fast Fourier and continuous wavelet transformation, bandpass filtering, and short-time Fourier transform using Matlab’s *spectrogram* function) and six connectivity measures (phase-locking value, Gaussian-Copula mutual information, Rayleigh test, weighted pairwise phase consistency, magnitude squared coherence, and entropy). We start our investigation by using simulated data where the connectivity between signals is precisely controlled. We then proceed to a single-subject real data set and finally to a full group analysis of an exemplary data set.

## 2. Material and methods

The evaluation of spectral estimation methods and connectivity measures is performed on two types of data. First, we use simulated data to control the type and extent of connectivity between the bivariate time series. Second, we use real MEG data from twenty participants listening to nine 1-min-long stories.

### 2.1 Data simulation

The simulated data is constructed by applying a fourth-order Butterworth bandpass filter (3-6 Hz) to a 1-minute simulated white noise signal (sampling rate: 100 Hz) with a mean of 0 and a standard deviation of 1. Two time series are then constructed by adding white noise (independently for each time series and with a mean of 0 and a standard deviation of 1) to the filtered noise. Therefore, the resulting time series show linear dependencies in the frequency range between 3-6 Hz that are evident as phase synchronisation and amplitude correlation. The degree of coupling can be adjusted through the amplitude of the added noise (see dedicated analyses below).

In what follows, the dependency between the time series will be quantified by applying all combinations of the six spectral estimation methods and the six undirected connectivity measures, which will be described in detail next.

### 2.2 Real data

We used MEG data recorded with a 275 whole-head sensor system (OMEGA 275, VSM Medtech Ltd., Vancouver, Canada) at a sampling frequency of 1200 Hz. The study was approved by the ethics committee of the University of Münster and conducted in accordance with the Declaration of Helsinki. Written informed consent was obtained before the measurement and participants received monetary compensation after the experiment.

Twenty native German-speaking participants (11 males, mean age 24.9 ± 2.6 years, range 20–32 years) listened to nine 1-min long audio recordings of their own voice in which they answered general questions such as ‘What does a typical weekend look like for you?’. Speech data was captured at a sampling rate of 44.1 kHz using a microphone placed at a distance of 155 cm from the participant’s mouth. Prior to data analysis, MEG data were visually inspected. No jump artifacts or bad channels were detected. A discrete Fourier transform (DFT) filter was applied to eliminate 50 Hz line noise from the continuous MEG data.

The wideband amplitude envelope of the speech signal was computed using the method presented in (Chandrasekaran et al., 2009). Nine logarithmically spaced frequency bands between 100-10000 Hz were constructed by bandpass filtering (third-order Butterworth filters). Then we computed the amplitude envelope for each frequency band as the absolute value of the Hilbert transform and downsampled them to 1200 Hz. We averaged them across bands and used the computed wideband amplitude envelope for all further analysis. Finally, MEG and speech envelope were downsampled to 256 Hz. In the preprocessing and data analysis steps, custom-made scripts in Matlab R2020 (The Mathworks, Natick, MA, USA) in combination with the Matlab-based FieldTrip toolbox (Oostenveld et al., 2011) were used following current MEG guidelines (Gross et al., 2013a).

For source localisation we aligned individual T1-weighted anatomical MRI scans with the digitized head shapes using the iterative closest point algorithm. Then, we segmented the MRI scans and generated single-shell volume conductor models (Nolte, 2003), and used this to create forward models. Next, the linearly constrained minimum variance (LCMV) algorithm was used to compute time series of voxels taken from a parcel showing medium connectivity (*L_PFop* located within the left inferior parietal lobule) of the volumetric HCP brain atlas (Glasser et al., 2016). The parcel selection was not relevant for the purpose of this study (which was focused on methods differences given two time series) but we ensured that the parcel showed significant connectivity to the speech envelope. The final time series representing activity from L_PFop was the first component of a singular value decomposition (SVD) of time series from all dipoles in this parcel.

### 2.3 Spectral estimation

Six different methods are used to perform a complex-valued spectral transformation of the time series in the frequency band. All methods except the wavelet transform use a frequency resolution of 0.5 Hz. For the subsequent connectivity estimation and evaluation we focused on the frequency band between 1 and 10 Hz.

1. The first three methods use the Fast Fourier transform (FFT) based implementation in FieldTrip (Oostenveld et al., 2011). The first method uses Hanning tapers while the second and third methods use discrete prolate spheroidal sequences (DPSS) in a multi-taper approach with ±1 Hz and ±2 Hz smoothing, respectively. In all three cases a 2s window with 50% overlap is used.
2. This uses the continuous wavelet transform implemented in Matlab with Morlet wavelets (cwtfilterbank.m with wavelet parameters 3 and 20). It uses L1-normalization so that equal amplitude oscillatory components at different scales have equal magnitude in the spectral estimate. The matlab function wt.m performs the actual transformation into the frequency domain.
3. A series of bandpass filters (windowed sinc FIR filter) is applied with edge frequencies that are 1 Hz below and above the center frequency. The center frequency changes from 1-10 Hz in steps of 0.5 Hz. The Hilbert transform is then applied for each filtered signal to obtain the complex-valued spectral estimate.
4. This spectral estimate is computed from Matlab’s *spectrogram* function in analogy to method 1. It also uses a 2s window with 50% overlap.

It should be noted that the number of complex valued data points returned from these methods is very different. Methods 1-3 and 6 are based on the FFT and return about one spectrum per second. Methods 4 and 5 instead return one spectrum per data sample and therefore provide many more, albeit largely redundant, data points. This has implications for computation time (see Table 1).

**Table 1.**
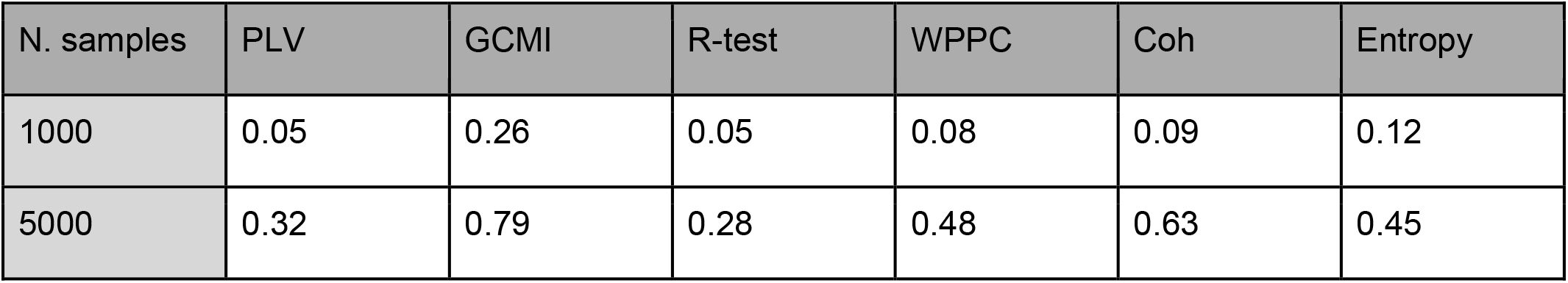
Computation time (in seconds) for different connectivity measures and two different numbers of samples in the complex frequency-domain input (3,8 GHz Quad-core i5 with 32 GB RAM). The mean over 100 repetitions is shown.

### 2.4 Connectivity measures

We use six undirected spectral connectivity measures:

1. Phase-locking value (plv; Lachaux et al., 1999): This is defined as the length of the vector average of the normalized (unit length) phase differences between time series x and y.
2. Gaussian-copula mutual information (gcmi; Ince et al., 2017): We compute mutual information between two bivariate time-series (real and imaginary part of x and y) using the original implementation (https://github.com/robince/gcmi).
3. Rayleigh test (R-test; Berens, 2009): The Rayleigh test is defined for circular (phase) data and tests for significant deviation from a uniform phase distribution. Here, it is applied to the phase difference.
4. Weighted pairwise phase consistency (wppc; Vinck et al., 2010): This measure does not directly test for a deviation of a phase distribution from a uniform distribution. Instead, it computes the pairwise difference of phases from this distribution. The rationale for this approach is that a preferred phase in the phase distribution would also lead to a cluster in the pairwise difference. However, in contrast to plv, wppc is not biased by the sample size. We compute wppc with code based on the FieldTrip implementation.
5. Magnitude squared coherence (coh): Coherence is a standard measure of association corresponding to a frequency domain correlation coefficient. It is computed by dividing the magnitude squared cross-spectral density between x and y by the product of the individual power spectra.
6. Entropy (ent; Shannon, 1948): We used entropy to quantify the deviation of the distribution of phase differences from a uniform distribution. In contrast to the other measures, this is sensitive to more than just unimodal phase difference distributions. Here, the computation uses a binning of phase differences into 20 bins.

### 2.5 Surrogate data and normalisation

For each connectivity measure, surrogate data are computed by randomly shifting the spectral estimates of one of the time series with respect to the other with a circular wrapping around the edges (using circshift.m in Matlab). This temporal shifting of data is an established technique for creating surrogate data because it destroys any true synchronisation in the data (Andrzejak et al., 2003) while preserving the signals’ autocorrelation structure. We perform this shifting procedure 200 times (unless otherwise stated) to create a distribution of 200 surrogate data points for each connectivity measure. Next, we normalise each connectivity measure by subtracting the mean and dividing by the standard deviation of the surrogate distribution for each frequency (Lancaster et al., 2018; Schreiber and Schmitz, 2000). This effectively normalises the connectivity measure and transforms it into units of standard deviations of the surrogate distribution. This useful normalisation makes measures more comparable to each other.

For our simulation, each combination of spectral estimation and connectivity measure is computed 500 times, with independently generated data in each iteration. Next, we define a performance measure D that quantifies the ‘average distance’ of the observed connectivity estimate from the 99th percentile of the surrogate distribution. This is computed as the mean of all connectivity values exceeding the 99th percentile of the surrogate distribution in the frequency band of simulated connectivity (3-6 Hz).

### 2.6 Data and code availability

We will make the Matlab code and underlying data publicly accessible in full through GitHub (https://github.com/IBiomag/).

## 3. Results

### 3.1 Comparison of combinations of spectral and connectivity estimates

First, we provide in Figure 1 an illustration of all combinations of spectral and connectivity measures for the simulated data described above (here with added noise with standard deviation of 1). For all of these combinations we plot the normalized connectivity spectrum (with the 95 percent bootstrap confidence interval) in the frequency range 0-10 Hz and the 99th percentile of the surrogate distribution (dashed line). All combinations of methods show a clear peak within the frequency band where connectivity was simulated (3-6 Hz). At the same time, it is clearly evident that results differ substantially in the shape of the spectrum and how far peaks are separated from the 99th percentile of the surrogate distribution (i.e., sensitivity for the true effect). First, for the same spectral estimate, different connectivity measures show markedly different sensitivity in detecting synchronisation in the data (compare panels within a row). That is, given the same information, the use of this information is significantly different between connectivity measures. Second, for the same connectivity measure, different spectral estimates lead to very different results (compare panels for a given column). Recall that synchronisation between time series x and y was simulated in the frequency band 3-6 Hz. Ideally, the spectrum in this band should exceed the 99th percentile line leading to a high D-value.

**Fig. 1.**
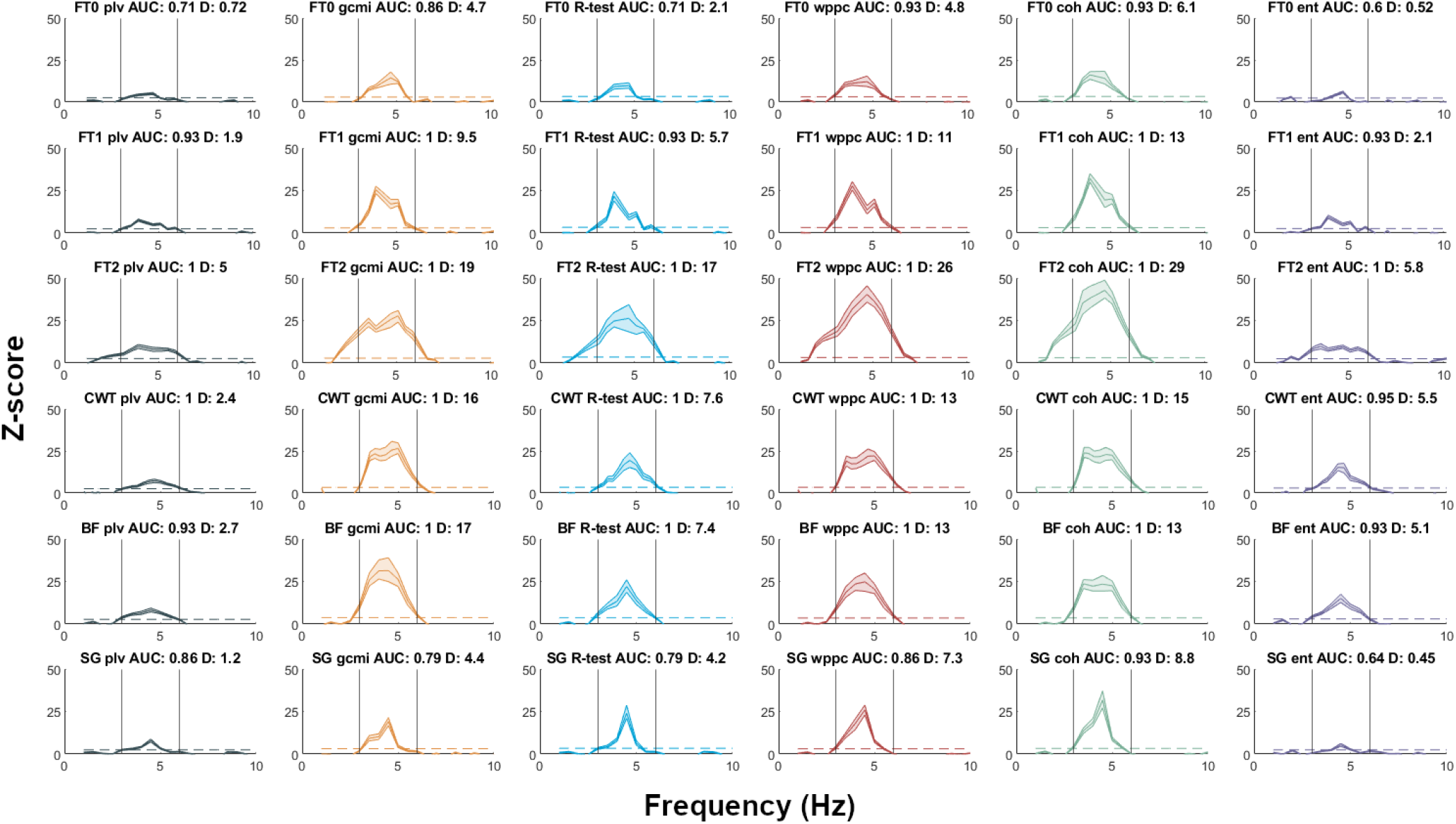
Connectivity spectra for all combinations of spectral estimates and connectivity measures. Connectivity was estimated for simulated data with a ground truth effect between 3-6 Hz (indicated by vertical lines) with an SNR of 1/20. The solid line shows the connectivity spectrum of a single trial z-scored with the mean and standard deviation of 200 time-shifted versions. The shaded area quantifies the uncertainty of the normalization and is based on the 95 percent bootstrap confidence interval of mean and standard deviation of the surrogate distribution. The dashed line represents the 99th percentile of the surrogate distribution. Each row is based on the same spectral estimate corresponding to the six methods in the same order as described in the methods section. Each column shows results from the same connectivity measure in the same order as described in the methods section. The title of each panel shows the spectral estimation method, the connectivity measure, the area under curve value (AUC), and the D-value defined in the methods section. FT0: FFT with Hanning taper; FT1: multitaper with ±1 Hz smoothing; FT2: multitaper with ±2 Hz smoothing; CWT: continuous wavelet transform; BF: bandpass filter; SG: spectrogram; plv: phase locking value; gcmi: gaussian copula mutual information; R-test: Rayleigh test; wppc: weighted pairwise phase consistency; coh: coherence; ent: entropy. The color code for connectivity measures is used throughout the manuscript.

From this simulation (based on 500 separate repetitions) we can already make several interesting observations. By comparing the different rows (spectral estimation methods), we note that the single taper FFT-based spectral estimates (FT0, SG) perform worse than the other methods (see Fig. 1, top and bottom row and note the individual scaling of each graph). An increased spectral smoothing with multitapers leads to an improved performance of all connectivity measures (higher D-values indicating larger separation from the surrogate distribution). However, this comes at the cost of a reduced spectral resolution which we will see in the analysis of real data (Fig. 6, third row from the top). Therefore, multitapers offer advantages for the detection of synchronisation (when the effect is not too narrow in the frequency domain) while they might be disadvantageous when trying to resolve different spectral peaks. Besides the FT2 method, the continuous wavelet transform, and bandpass filtering perform very well (Fig. 1, second and third row from the bottom).

A comparison of connectivity measures (different columns) reveals best performance for *wppc* (shown in red) followed by *gcmi* (orange). In contrast, *ent* (purple) and *plv* (grey) show relatively poor performance. Overall, simulation-based connectivity spectra suggest that the combination of FT2 and *wppc* shows the best performance.

In order to look at performance differences in more detail, we conducted pairwise comparisons of all 36 possible combinations (6 spectral estimates x 6 connectivity measures). Specifically, we computed Cohen’s d as a measure of effect size separating the D values from the 500 simulations of each combination (see Fig. 2a). Not counting the main diagonal of the symmetrical 36 × 36 matrix, we gained 35 effect sizes for each combination of spectral estimate and connectivity measure. The respective distributions are shown in Fig. 2b. Overall, pairwise comparisons corroborate the previous impression that wppc with FT2 outperformed most of the other combinations: Judging by the box plot notches in Fig. 2b, only gcmi (with FT2, CWT, or BF) and the R-test (with FT2) reached a similar performance. Moreover, the performance for entropy combined with FT0 or SG was particularly subpar, paralleled only by plv combined with the same estimates. Finally, pairwise comparisons supported the initial impression of lowered performance of FT0 and SG in all combinations, irrespective of the connectivity measure (see Fig. 2b).

**Fig. 2.**
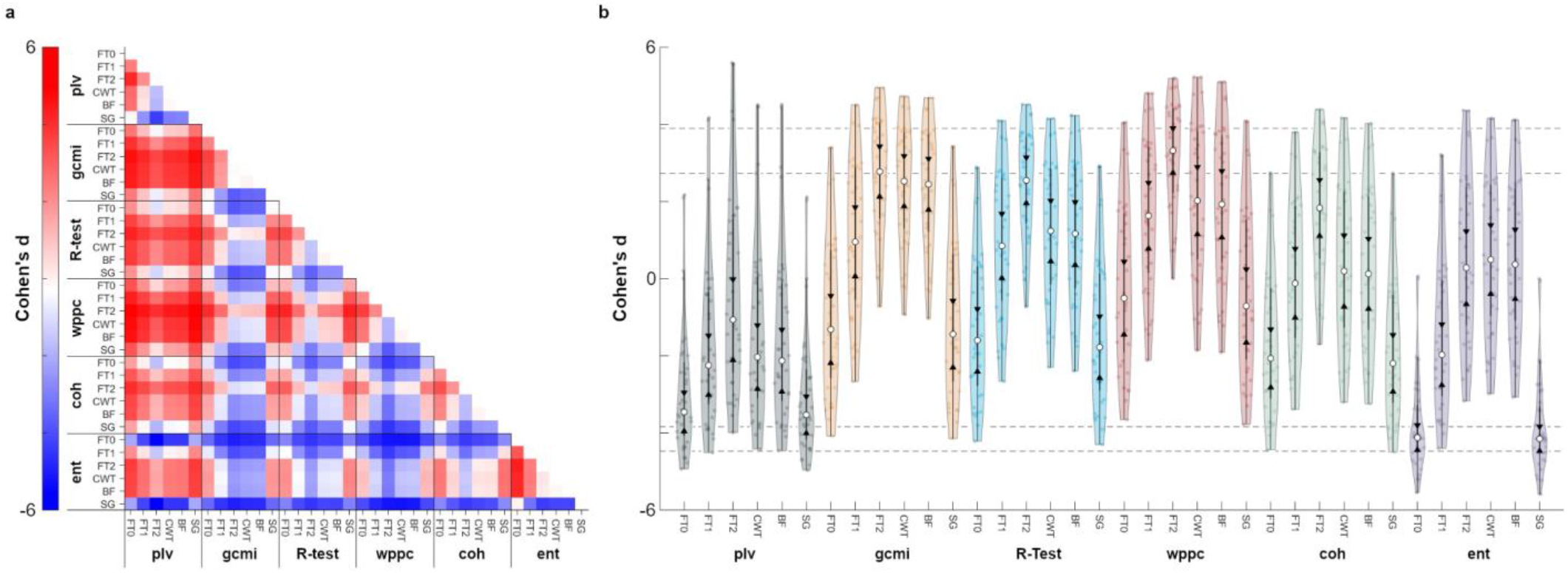
Pairwise comparisons of spectral estimates and connectivity measures. a, To assess performance differences within the simulated data, we compared each combination of spectral estimate and connectivity measure with any other combination, resulting in a 36 × 36 symmetrical matrix. We computed Cohen’s d as a measure of effect size separating the D-values from the respective 500 simulation iterations of any two combinations. Positive values indicate higher D values for the row (vs the column) combination. b, Violin plot shows the distribution of effect sizes for each of the 36 combinations (grouped according to connectivity measures). White dots mark the respective median of each combination, black triangles indicate box plot notches for comparison across combinations. As a reference, top dashed lines indicate box plot notches for wppc with FT2 estimation, which showed the best median performance overall. Similarly, bottom dashed lines indicate box plot notches for entropy with SG estimation whose performance was lowest overall.

### 3.2 Effect of SNR

Next, we aimed to quantify the effect of different levels of signal-to-noise ratio (SNR) on performance. This was motivated by the hypothesis that different connectivity measures are differentially sensitive to varying SNR levels. Indeed, this can be seen in Figure 3 which follows the arrangements of rows and columns from Figure 1. Towards the right of the figure, the SNR is increasing. A differential SNR-effect on performance is quite prominent in the comparison of the third and fifth column. While wppc (shown in red) is the most sensitive measure in the middle column (SNR parameter = 1.5) it is outperformed by gcmi (yellow) for the highest SNR (SNR parameter = 2.5, rightmost column). This indicates that performance of gcmi increases more strongly with SNR than for other measures. This high performance for high-SNR data was also described in the original gcmi publication (Ince et al., 2017). While all measures benefit to some extent from SNR-increases (albeit none as much as gcmi), this benefit is considerably lower for plv (grey) and entropy (purple) compared to the other measures. Interestingly, the SNR-dependence of performance increase is rather similar across spectral estimation methods (e.g. the order of connectivity measures according to performance in the rightmost column is almost identical across spectral estimation methods (rows)). Still, the absolute D-values are very different across rows and show best performance for FT2 and BF and, as before, worst performance for FT0 and SG.

**Fig. 3.**
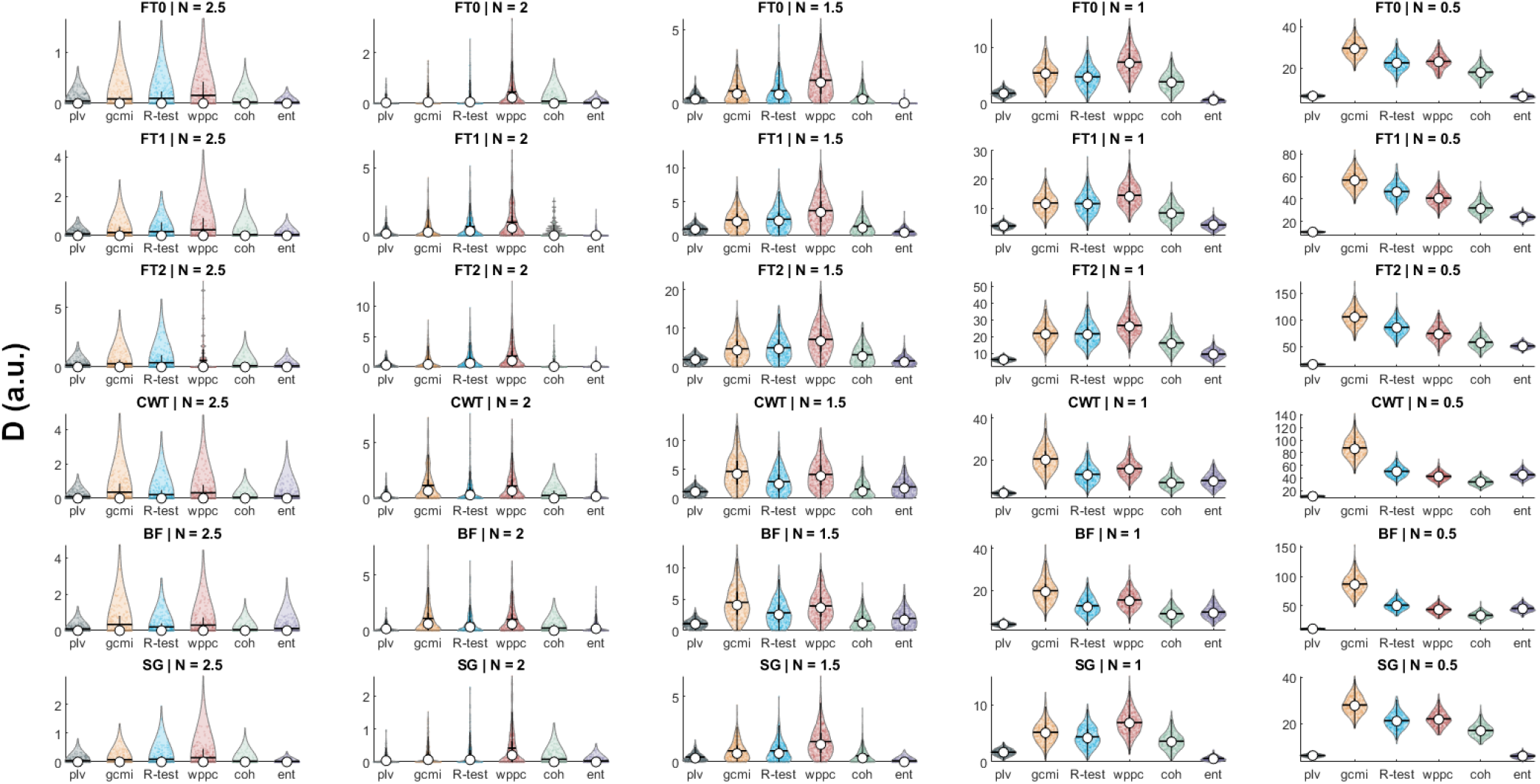
Effect of SNR. Each subplot shows a violin plot for each of the six connectivity measures (same order as in Fig. 1) of the D-value across 500 repetitions of the simulation. Columns correspond to different SNRs. The noise factor (N) specifies the standard deviation of the noise added to the signal. SNR increases from left to right.

### 3.3 Effect of downsampling spectral estimate

Performing spectral estimation with either bandpass filtering and Hilbert transformation or the continuous wavelet transform leads to many more samples compared to FFT-based methods. This results in longer computation times for these continuous methods when computing connectivity measures (see Table 1). Especially at low frequencies, the continuous spectral estimates show substantial redundancies between neighbouring samples. Therefore, we investigated the effect of downsampling the continuous spectral estimate by a factor of 10 on the sensitivity of the connectivity measure. Figure 3 shows violin plots of the distribution of D-values across 500 iterations of our simulation. For each connectivity measure the darker color (left plot of each pair) shows the original result and the lighter color (right) shows the result from the downsampled spectral estimate. As can be seen, results are very similar for original and downsampled spectral estimates for all connectivity measures. A linear mixed effects model (LMEM) indicates a significant effect of downsampling (β = - 0.09, t(11992) = -2.25; p = .024, D = *β*_0_ + *β*_1_ * spec + *β*_2_ * conn +*β*_3_ * ds + e_j_ ; spec, conn, ds are categorical variables for spectral estimation method, connectivity method and downsampling, respectively). However, the rather small LMEM estimate of the change in D-value with downsampling makes it negligible for practical applications. This indicates that, for the frequencies considered here, results are not much affected by downsampling while computation time decreases (see Fig. 4).

**Fig. 4.**
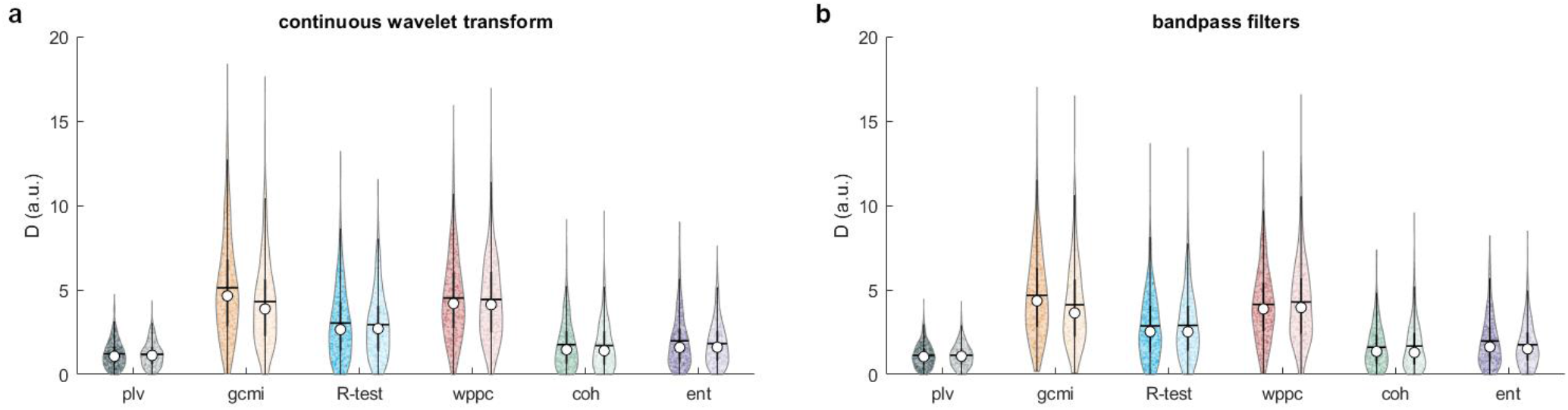
Effect of downsampling on the two continuous spectral estimates, continuous wavelet transform (a) and bandpass filter (b). For each connectivity measure, two violin plots show the distribution of D-values for 500 repetitions of the simulation for the original (sampling frequency = 100 Hz, darker colours) and the downsampled spectral estimate (sampling frequency = 10Hz, lighter colours).

### 3.4 Deviation from unimodal phase distribution

Ideally, connectivity measures should be sensitive to any deviation of the phase distribution from a uniform distribution. Here, we test the specific case of a bimodal phase distribution. For the first half of the time series we simulate a zero-degree phase synchronization while the second half uses a simulation of a 180-degree phase difference between both signals. This results in a bimodal phase distribution with deviation from a uniform distribution at opposite sides of the circular phase space. Clearly, all connectivity measures except entropy (shown in purple) fail to capture this more complex phase dependency (see Fig. 5). Given the definition of these measures, this result is not surprising: In all measures (except entropy) the opposite phase differences across the unit circle lead to cancellation and result in a non-detectable phase synchronization. Entropy instead quantifies any deviation from a uniform distribution in phase bins across the unit circle and therefore captures this bimodal phase distribution. However, as we could see from the previous section, this sensitivity to more complex deviations from a uniform distribution leads to a reduced sensitivity for unimodal phase distributions (see Fig. 1 and 2).

**Fig. 5.**
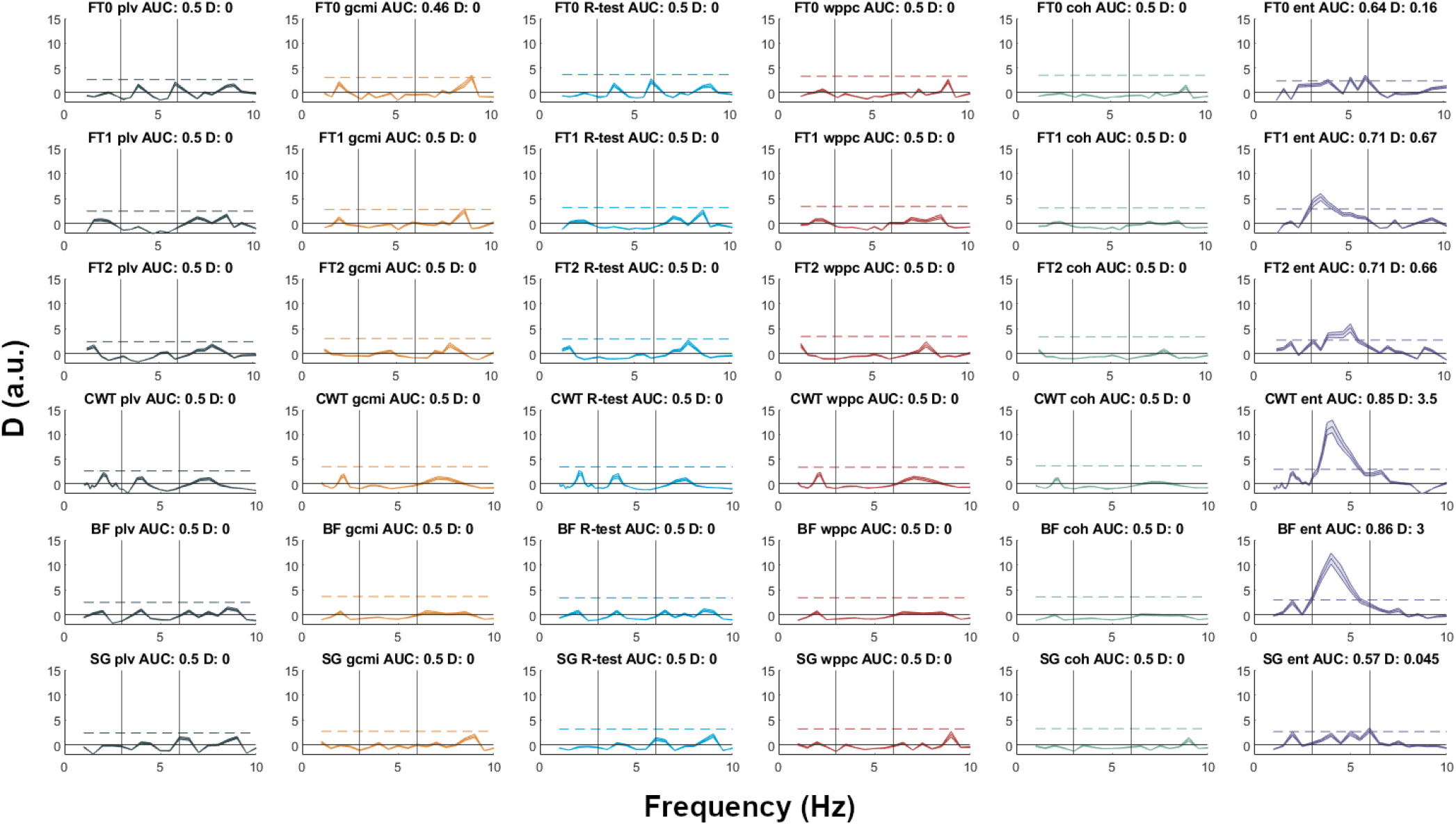
Deviation from unimodal phase difference distribution. The layout is the same as in Figure 1. The underlying data lead to a bimodal phase distribution that is only detected by the entropy measure.

**Fig. 6.**
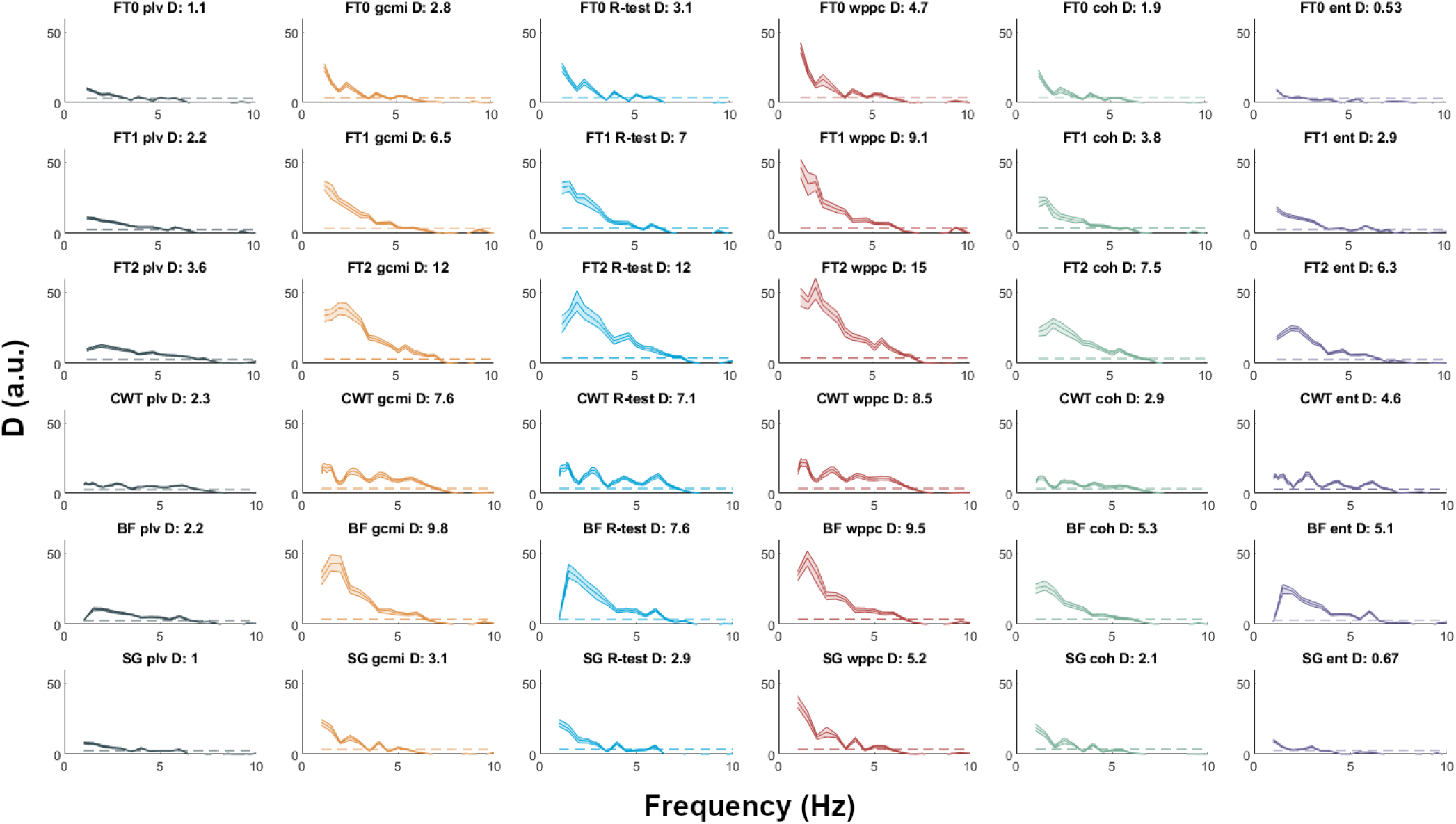
Results for 9-min long MEG recording. Layout is identical to Figure 1.

### 3.5 Real data

Next, we compared the same combinations of spectral estimation and connectivity methods in real data. Before proceeding to group analysis, we studied speech envelope to MEG connectivity spectra in a single 9-min long data set. Figure 6 shows the results following the same computations and plotting format as in our simulated data. Results are generally consistent with our findings from simulated data (see Fig. 1). Overall, best performance can be seen for FT2 and wppc (third row from the top, red) followed by gcmi (yellow) and Rayleigh test (blue). Interestingly, this computation on real data shows that the spectral structure is mostly determined by the spectral estimate and not so much by the connectivity method (i.e., spectra in a row are more similar than spectra in a column). Obviously, there is more spectral structure in real data than in the simulated data where only a single spectral peak was evident. Not surprisingly, this spectral structure is mostly lost in FT2 due to the spectral smoothing of +/-2Hz. Instead, the highest complexity of spectral structure can be seen using the continuous wavelet transform (CWT, third row from the bottom) and still leads to high sensitivity (large D-values) compared to FT2. CWT is therefore probably most appropriate when preservation of the spectral structure is important for the research question at hand. However, the ‘true’ spectral structure of the data is unknown so we cannot evaluate and compare the performance of spectral estimation measures in this regard.

### 3.6 Data length and computation time

The dependence of statistical effect sizes on data length for different combinations of spectral estimation and connectivity measure is of considerable practical importance. An optimal combination can lead to increased statistical sensitivity in shorter recordings. Figure 7 shows the dependence of D-values on data length. Each subpanel shows results for the six spectral estimation methods in the order used in all other plots (FT0, FT1, FT2, CWT, BF, SG). Each subpanel has six groups of bar plots corresponding to the six connectivity measures (plv, gcmi, R-test, wppc, coh, ent) and each group of bar plots shows the D-values for nine linearly spaced data lengths from 1-9 mins. As expected, D-values increase in general with increasing data length and in most cases even from 8 min to 9 min. Our results also illustrate that the combination of methods clearly matters. For example, using FT0 and PLV (top left, grey) for 9 min data leads to worse performance than FT2 and wppc (top right, red) for 2 min data (at least for our definition of performance and our implementation of methods).

**Fig. 7.**
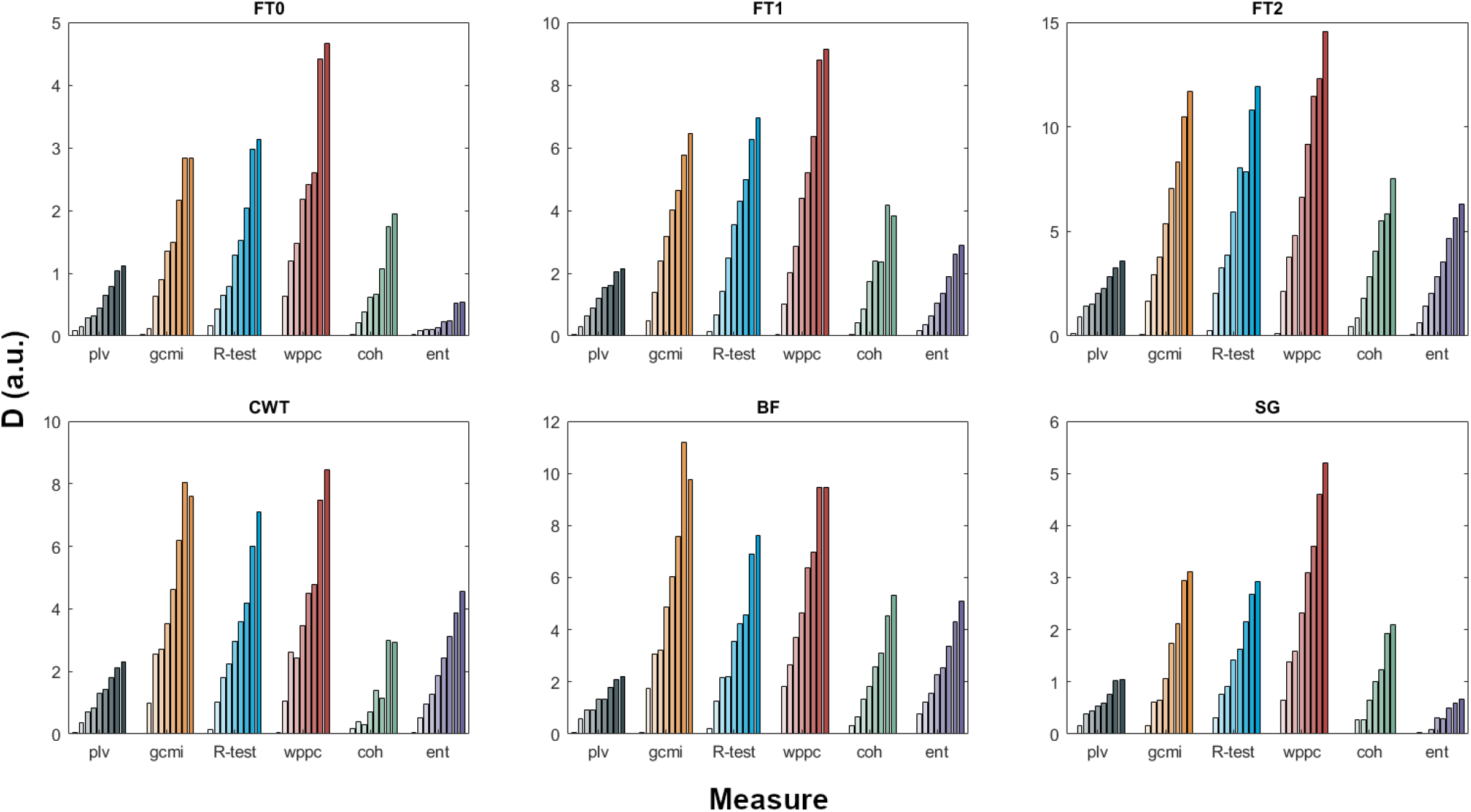
Effect of data length. Each subplot corresponds to one spectral estimation method. In each subplot colored bars show the nine D-values for data length from 1-9 minutes for each of the six connectivity measures.

Another point of potential practical importance is computation time. Table 1 compares computation time (including 200 surrogate computations) for the six different connectivity measures and two different numbers of samples in the input for our implementation of the methods, based on our implementation. Computation times are all in a similar range while gcmi is the slowest method and plv and R-test the fastest. The exact times of course depend on the computer architecture and we show this table mainly to allow comparison across methods. If computation time is a major concern, then R-test should be preferred over plv given its superior performance in all our results (both simulated and real data).

### 3.7 Effect of number of surrogate computations

Our measure of performance, D-value, is derived from a distribution of surrogate data (see Methods section). Here we address the question to what extent D depends on the number of surrogate data realisations. Figure 8 follows the layout of Figure 7 and shows D for 9 min of data for three different numbers of surrogate data (100: left bar; 200: middle bar; 400 right bar). Interestingly, D-value changes very little for different numbers of surrogate data realisations. However, we would like to note that the bootstrap confidence interval (shown as shaded area for example in Fig. 1) decreases with increasing number of surrogate data realisations. For practical applications, 100 or 200 surrogates seem to be sufficient, as the incremental change in D for more surrogate iterations is negligible.

**Fig. 8.**
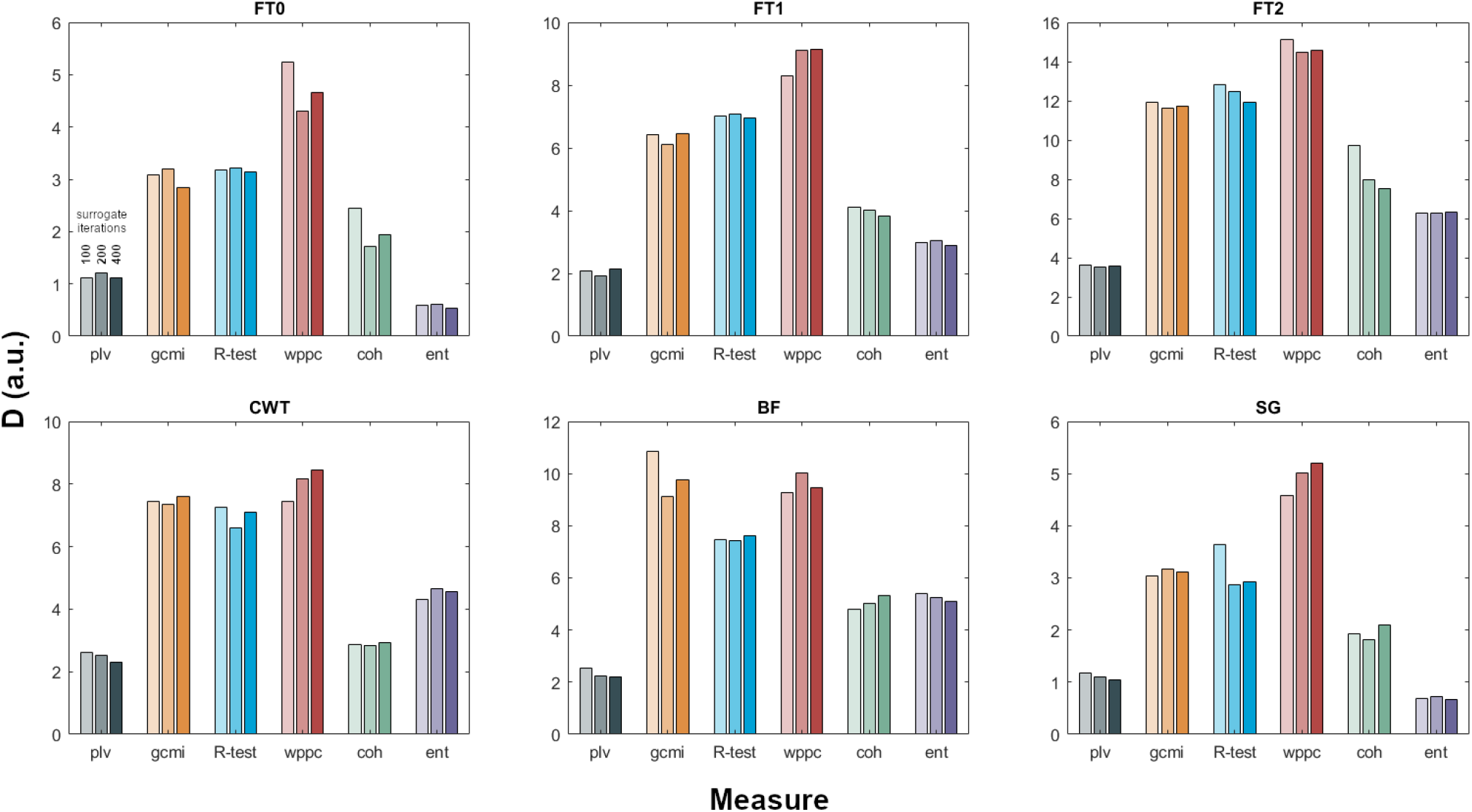
Effect of number of surrogate data on D for 100 (left bar), 200 (middle bar) and 400 (right bar) surrogates.

### 3.8 Group statistics

In the previous sections we have exclusively used single simulated or real data sets to compare performance of different spectral estimation and connectivity techniques. In our final analysis we will now extend this approach to group analysis. For data from 20 participants we repeated the computations shown in Fig. 6, resulting in normalised connectivity spectra. We then performed standard group analysis using independent samples t-test against a fixed value of 1.64 corresponding to the 95th percentile of a normal distribution. Statistical significance was established with non-parametric cluster-based permutation tests as implemented in FieldTrip with 2000 randomizations.

Figure 9 shows spectra of t-values for the different combinations of spectral estimates and connectivity measures. First, comparing spectral estimates we find that the multi-taper spectral estimate with smoothing of +/-2Hz (third row from the top) performs best, followed by the bandpass filter (second row from the bottom). The comparison of connectivity measures (different columns) shows markedly smaller differences in group results than in the single data sets. Surprisingly, plv (grey) performs much better in group statistics compared to the single simulated and real data sets. Overall, in our group analysis, the choice of spectral estimation method appears to be more important than the connectivity measure.

**Fig. 9.**
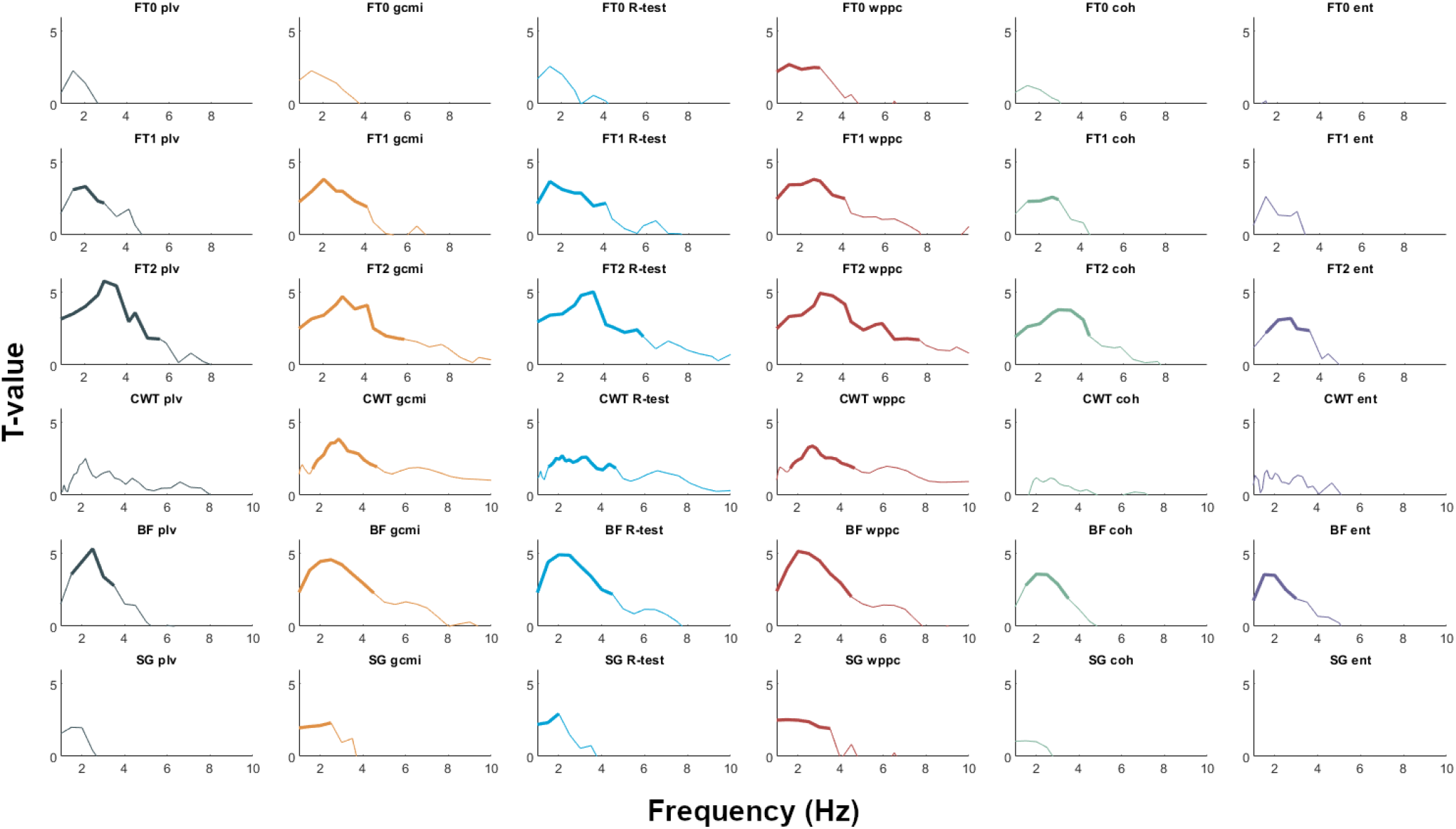
Group statistics. Layout is identical to Figures 1 and 6. T-values are plotted between 1-10 Hz. Cluster-corrected significant frequency bands are marked by increased line width.

## 4. Discussion

In this study we aimed to demonstrate how the sensitivity to detect cerebro-peripheral connectivity is affected by different combinations of spectral estimates and connectivity measures. Results from simulated and real data reveal conclusively that the selection of methods can facilitate or preclude the detection of significant connectivity, both at the individual and the group level.

Spectral estimates and connectivity measures interact with each other in non-trivial ways. For a given spectral estimate the available information about the underlying synchrony is utilized by different connectivity measures in markedly different ways. More precisely, if phase synchronization exists in the data (as in our simulated data) the distance of estimated connectivity from the surrogate distribution varies considerably across connectivity measures.

Regarding spectral estimation methods, we compared different Fourier-based techniques that mostly differ in their spectral smoothing, with wavelet spectral estimates and those based on bandpass filtering followed by Hilbert transformation. Overall, highest performance was observed for FT2, the multi-taper approach with +/-2 Hz spectral smoothing. CWT and BF performed also well and in general better than FT0 and SG. Conceptually, Fourier-based methods, Hilbert transformation, and wavelet transformation are very different, but it has been shown that - given well-chosen parameter settings - these three approaches can lead to converging results (Bruns, 2004). In our analysis, we used implementations with standard parameter settings. This might in part explain the difference in performance between FT0 and SG on the one hand, and between CWT and BF on the other hand. Both FT0 and SG reflect overlapping 2-second window FFT-based estimates, with a single Hanning taper applied to each data window. In the simulations, this resulted in 59 degrees-of-freedom for the spectral amplitude and phase estimates, one for each window. In comparison, both CWT and BF resulted in a single amplitude and phase estimate per original time point, which, even considering the large amount of redundancy for consecutive time points, likely led to more stable estimates. Multi-taper based spectral estimation (Percival and Walden, 1993) trades spectral resolution for reduced variance in the spectral estimates, thus increasing sensitivity. This is also referred to as spectral smoothing, and is achieved by applying a set of tapers to the data, the number of which is determined by the time-bandwidth product NW, i.e. the length of the data segments (N) multiplied by the specified smoothing parameter (W). The number of tapers used is then typically 2NW-1. In our case, as both FT1 and FT2 were implemented using 2-second long overlapping data windows, the smoothing increased the degrees-of-freedom for the spectral estimates by a factor of 3 and 7, respectively. In general, we can expect that an analysis is optimal when the effective resolution of its spectral estimate is adjusted to the expected bandwidth of significant phase synchronization (which is unknown in real data). For example, if phase synchronization exists in a 4 Hz wide frequency band (e.g. 8-12 Hz) then multi-taper smoothing of +/-2 Hz should be optimal. This is largely what we observe here. However, additional factors apparently contribute to performance. For example, our simulation contained significant synchronization over a 3 Hz bandwidth. Therefore, if spectral smoothing were the only factor determining analysis performance then we would expect the +/-1 Hz and +/-2 Hz smoothing to perform equally well. The fact that +/-2 Hz multitaper analysis performs better than other spectral estimates with less or no spectral smoothing indicates that the smoothing itself improves analysis sensitivity, albeit at the cost of reduced spectral resolution. Spectral resolution should be highest for CWT where different wavelets capture spectral structure even at low frequencies. Indeed, this point is nicely illustrated in Fig. 6. Whereas CWT-based connectivity spectra show separate peaks at low frequencies, these are largely merged into one for the +/-2 Hz multitaper estimate. Since in real data the underlying spectral structure is unknown it might be advisable to use two approaches, the FT2 computation for optimal sensitivity and CWT for optimal spectral resolution. Alternatively, longer data segments can be defined for the spectral transformation, which would then still allow for leveraging increased sensitivity of the multi-taper framework. For instance, increasing the window length from 2 seconds to 4 seconds would allow for a reduction of the smoothing parameter from 2 to 1 without compromising the number of tapers applied.

We non-exhaustively compared six different connectivity metrics aimed at capturing band-limited phase synchronization between signals. In most cases the weighted pairwise phase consistency (wppc) outperformed the other methods. The main exception was the improved performance of Gaussian copula based mutual information (gcmi) for data with high SNR. In general, gcmi and R-test performed also very well. Performance for coherence (coh) was overall quite good (particularly in the simulations), and performance for phase locking value (plv) and entropy (ent) was lowest overall. The entropy measure, however, was the only metric that proved sensitive to more complex distributions of phase differences. Here, we tested the challenging case of a bimodal distribution of phase differences, with the modes of the distribution 180 degrees apart, that leads to cancellation in most methods and a failure to detect this more complex phase synchronization.

(Weighted) ppc (Vinck et al., 2010) has been proposed as a metric that provides a bias-free estimate of phase synchronisation, as opposed to the more traditionally used phase locking value or coherence coefficient. Its improved performance could result from this reduced bias, possibly due to a reduction in variance of the surrogate distribution, as well as a shift towards zero. Our implementation of gcmi used both amplitude and phase information for the estimation of the connectivity, just like wppc and coh. R-test, plv, and entropy only use the phase information. Obviously, the sensitivity of a particular metric is in part determined by the actual functional statistical relationship between the measured signals. If the relationship is mainly expressed in terms of the phase difference, then ‘phase only’ metrics will be sufficient. If the relationship is in part also expressed in terms of the amplitude correlations, then ‘phase and amplitude’ metrics will be more sensitive. Non-linear relationships might be more easily captured with gcmi or entropy.

Another point of practical importance for the design of cerebro-peripheral connectivity studies is the required data length. We compared performance of different combinations of spectral estimates and connectivity measures for data length between 1-9 min. In almost all cases, the mean distance of estimated connectivity relative to the surrogate distribution increased continuously with increasing data length. Therefore, statistical analysis will benefit from long recordings (see e.g. Daube et al., 2019), particularly if subtle experimental effects are to be detected.

In summary, our analysis of cerebro-peripheral connectivity has revealed that results depend significantly on the combination of spectral estimation and connectivity measures. Our analysis of simulated and real data provides some observations that might assist scientists in this field in making a more informed choice of analysis methods given their respective priorities. We hope that this leads to further advances in the exciting field of cerebro-peripheral connectivity analysis.

## Acknowledgements

This work was supported by the Interdisciplinary Center for Clinical Research (IZKF) of the medical faculty of Münster (Gro3/001/19) and the DFG (GR 2024/5-1). This work was further supported by The Netherlands Organisation for Scientific Research (NWO Vidi: 864.14.011) to JMS. The authors would like to thank Karin Wilken, Ute Trompeter, and Hildegard Deitermann for their invaluable assistance during data collection.

## Supplemental Figure

**Fig. S1.**
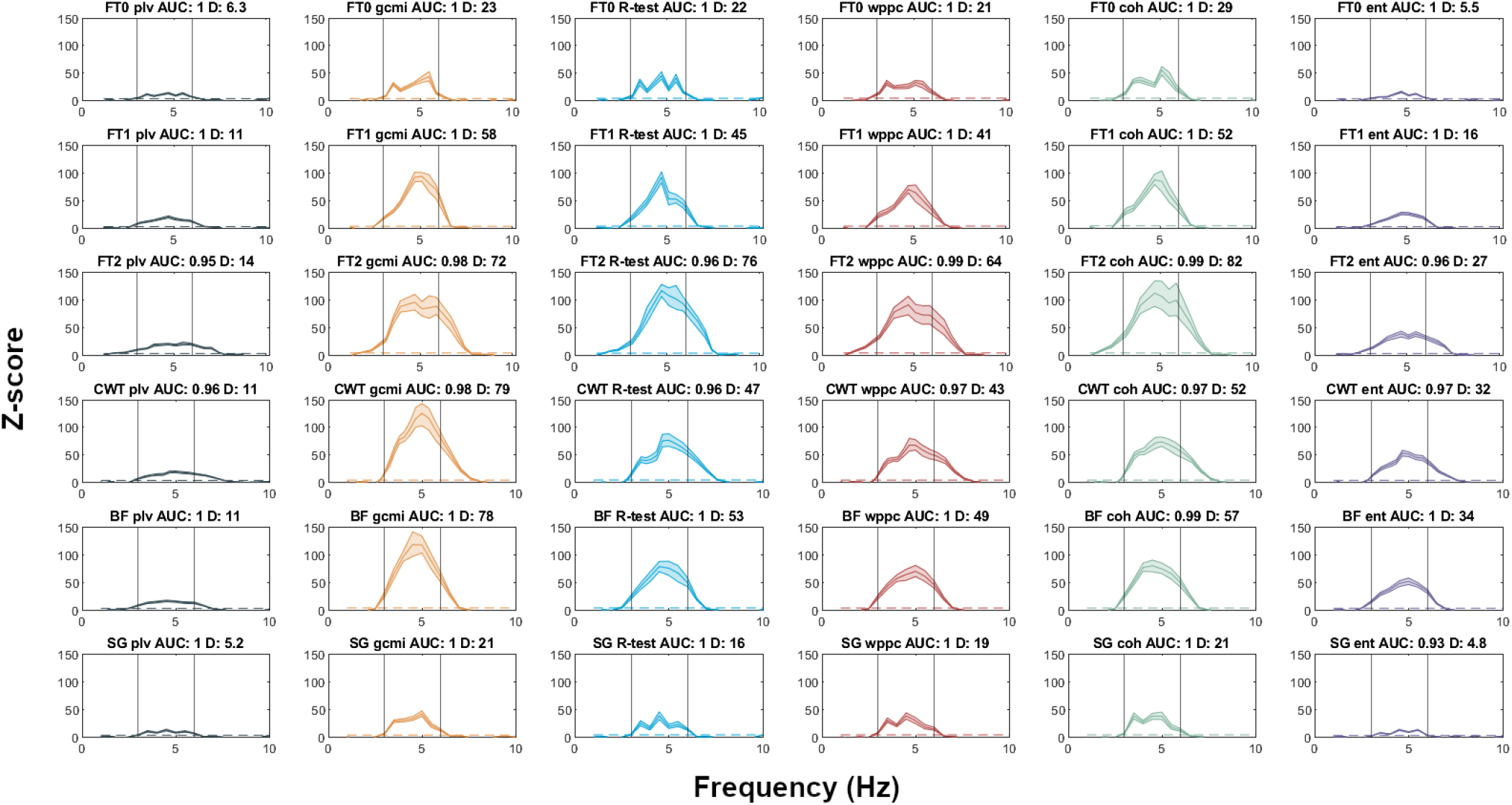
Same as Figure 1, but simulated with 1/f noise.

